# PHYSICAL EXERCISE RESTORES NEUROCOGNITIVE HOMEOSTASIS DISRUPTED BY NON-SEVERE MURINE MALARIA

**DOI:** 10.1101/2025.01.08.631885

**Authors:** Luciana Pereira de Sousa, Ingrid de Oliveira Lavigne, Mônica da Silva Nogueira, Cláudio Tadeu Daniel-Ribeiro

**Affiliations:** Laboratório de Pesquisa em Malária, Instituto Oswaldo Cruz (IOC), Fundação Oswaldo Cruz (Fiocruz), Rio de Janeiro, RJ, Brazil; Centro de Experimentação Animal, IOC, Fiocruz, Ministério da Saúde, Rio de Janeiro, RJ, Brazil; Centro de Pesquisa, Diagnóstico e Treinamento em Malária (CPD-Mal), Fiocruz and Secretaria de Vigilância em Saúde e Ambiente, Ministério da Saúde, Rio de Janeiro, RJ, Brazil

## Abstract

Malaria disrupts neurocognitive homeostasis in humans, including in the non-severe manifestation of the disease - which is the most prevalent form of malaria in the world. This disruption is classically observed in human and experimental models of cerebral malaria. More recently, we demonstrated that this can also be observed in an experimental model of non-severe malaria and that Th2-immune response improves cognition and attenuates anxiety-like behavior associated to malaria. Complementarily, we have been studying the effect of physical exercise in restoring the neurocognitive homeostasis lost after non-severe murine malaria.

## INTRODUCTION

Malaria is associated with neurocognitive sequelae in humans and mice, especially in its most severe form, cerebral malaria (CM) (WHO, 2024; Reis et al., 2012; for review see Rosa-Gonçalves et al., 2022; de Sousa et al., 2023). In non-severe malaria (n-SM), cognitive deficits related to learning and memory are also observed in humans, affecting the well-being of children during neurodevelopmental and school stages, as well as the elderly and even asymptomatic individuals (Fernando et al., 2003; Vitor-Silva et al., 2009; Fink et al., 2013; Nankabirwa et al., 2013; Tapajos et al., 2019; Pessoa et al., 2022;). Considering the strong interaction between the plastic cognitive immune and nervous systems (Cohen 1992; Kipnis 2016; de Sousa et al., 2023), we studied the effect of Th1 and Th2 immune response inducing stimuli on neurocognitive performance in healthy C57BL/6 mice and in the attenuation of neurocognitive sequelae recorded after a single episode of *Plasmodium berghei* ANKA (*Pb*A) n-SM treated before progression to CM (de Sousa et al., 2018). A positive effect of Th2-immune stimulation improving cognition in homeostasis and mitigating neurocognitive sequelae was demonstrated (de Sousa et al., 2021), identifying that only one of the components of the Th2-immunostimulatory strategy, the tetanus-diphtheria vaccine (Td), couldrestore the neurocognitive normality impacted by n-SM (Rosa-Gonçalves et al., 2022).

Knowing that physical exercise (PE) may also influence the activity of the immune system and is a beneficial promoter of cognition (Seo et al., 2019; Scheffer et al., 2020; Pahlavani et al., 2023), the objective of this work is to investigate the effect of PE, isolated or in combination with immunization with Td vaccine, in promoting cognitive-behavioral performance in homeostasis and mitigating neurocognitive sequelae resulting from experimental murine n-SM.

## METHODOLOGY

### Animals

The study was approved by the Ethics Committee for Scientific Use of Animals of the Instituto Oswaldo Cruz of Fiocruz (CEUA/IOC, L-004/2020-A2). Seven-week-old C57BL/6 female mice, weighing 20–25g, were provided by the *Instituto de Ciência e Tecnologia em BioModelos* of the *Fundação Oswaldo Cruz* (*ICTB-Fiocruz*, Brazil). Mice were kept in groups of five per cage, housed in racks with an air filtration system, in a room maintained at 25 °C with 12-hour light/dark cycles with free access to food and water.

### Infection and treatment

C57BL/6 mice were divided into groups infected or not with 10^6^ *Plasmodium berghei* ANKA (*Pb*A)-parasitized erythrocytes per 100 μl inoculum, and injected by the intraperitoneal (ip) route, treated (both infected and control groups) on the 4th day after infection with 25 mg/Kg of chloroquine by gavage for seven days (de Sousa et al., 2018).

### Physical exercise and immunization

mice were either subjected to PE practice or not at 14 (immediate intervention) or 80 (delayed intervention) days after the end of antimalarial therapy. A subset of each group was also subjected to immunization with Td-vaccine or not at 14 days after the end of antimalarial therapy, being three doses every twenty days. PE consisted of a moderate aerobic involuntary practice performed on a motorized treadmill for mice in different protocols (de Sousa et al., 2021), for one or two months in five or three weekly sessions, respectively, at speeds of 8, 12 and 8m/min for 5, 20 and 5min, respectively (Figure 1). For the practice of immediate intervention, the protocol of PE three times a week for two months was considered, in combination or not with stimulation of the immune system. For the delayed intervention group, we applied PE five times a week for a month, as part of a pilot study (a comparison between the two protocols is being conducted).

**Figure 1.**
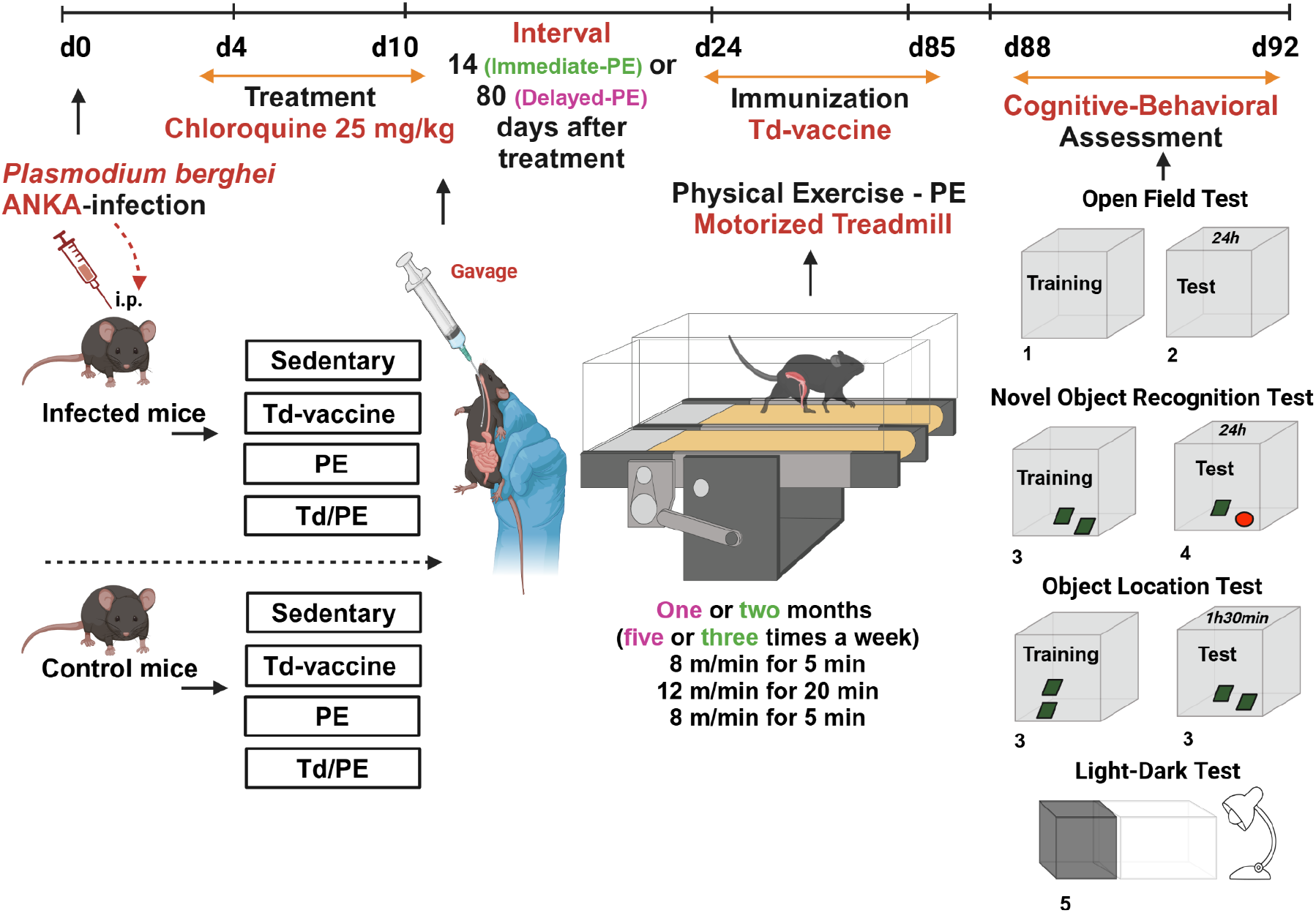
Flowchart of experiments. C57BL/6 mice are infected or not with *Plasmodium berghei* ANKA and treated on the fourth day after infection with 25 mg/kg of chloroquine via gavage for seven days, then subdivided into groups of sedentary, sedentary and immunized with dT vaccine, exercised, and exercised and immunized with dT vaccine. Immunization and exercise were started 14 days after the end of treatment. Delayed-PE was alternatively initiated 80 days after the end of treatment. Finally, the animals had their cognitive and behavioral performance evaluated in tasks such as open field, novel object recognition and light-dark, from day 88 on, to evaluate parameters such as locomotion, long-term memory, anxiety-like behavior, and object location test, as previously described (de Sousa et al., 2018; Vogel-Ciernia & Wood, 2014).

### Behavioural tasks

Animals were then subjected to tasks such as open field (to analyze locomotion and habituation memory), novel object recognition (to test long-term memory) and light-dark (to measure anxiety-like behavior), and object location test (to measure short-term memory), as previously described (de Sousa et al., 2018; de Sousa et al., 2021; Rosa-Gonçalves et al., 2022; Vogel-Ciernia & Wood, 2014) (Figure 1).

### Data analyses

The data were acquired and analyzed by the any-maze tracking software (Stoelting Co, USA, 7.4 version). Statistical analyses of the data were performed using GraphPad Prism by Dotmatics software (version 8.0).

## RESULTS

Infected/sedentary/not immunized mice group (n. = 11) showed cognitive deficits in long-term memory and anxiety-like behavior when compared to the control - healthy/sedentary/not immunized - group (n. = 11) (*P < 0*.*0001\*\*\** and *0*.*0015\*\**, respectively), in accordance with what is traditionally observed by our group (Figure 2). Immediate PE was more effective in attenuating neurocognitive damage of n-SM than the Delayed-PE. The infected/exercised/immunized-or not mice groups (n. = 11 for each group) presented better results, in relation to the infected/sedentary/not immunized group (*P<0*.*0001***, for both comparisons*), which were similar to the control group (*P = 0*.*0960; 0*.*3372; 0*.*9470*) (Figure 2).

**Figure 2.**
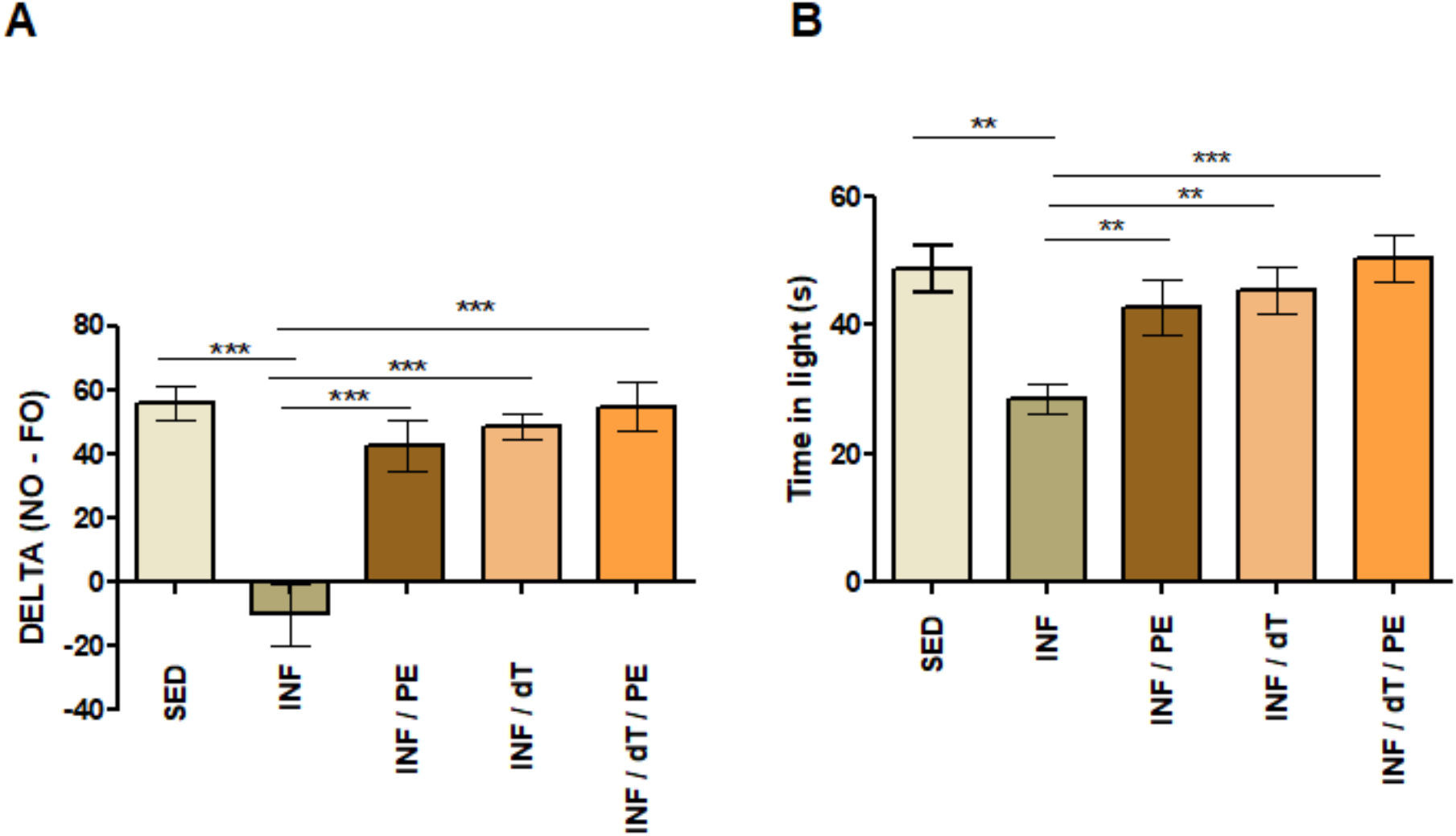
Effect of immediate physical exercise and immunization with Td vaccine. started 14 days after the end of chloroquine treatment - in mitigating cognitive (A) and behavioral (B) deficits caused by non-severe murine malaria.

Considering animals in homeostasis, PE, whether combined with immunization or not (n. = 10 for each group), is capable of improving cognitive performance when compared to the control - healthy/sedentary - group (*P = 0*.*0424**; 0*.*0071\*\**), as well as improving the natural behavior of the animals regarding the default “anxiety-like” pattern (*P = 0*.*0207**; 0*.*0134\*\*\**) (Figure 3).

**Figure 3.**
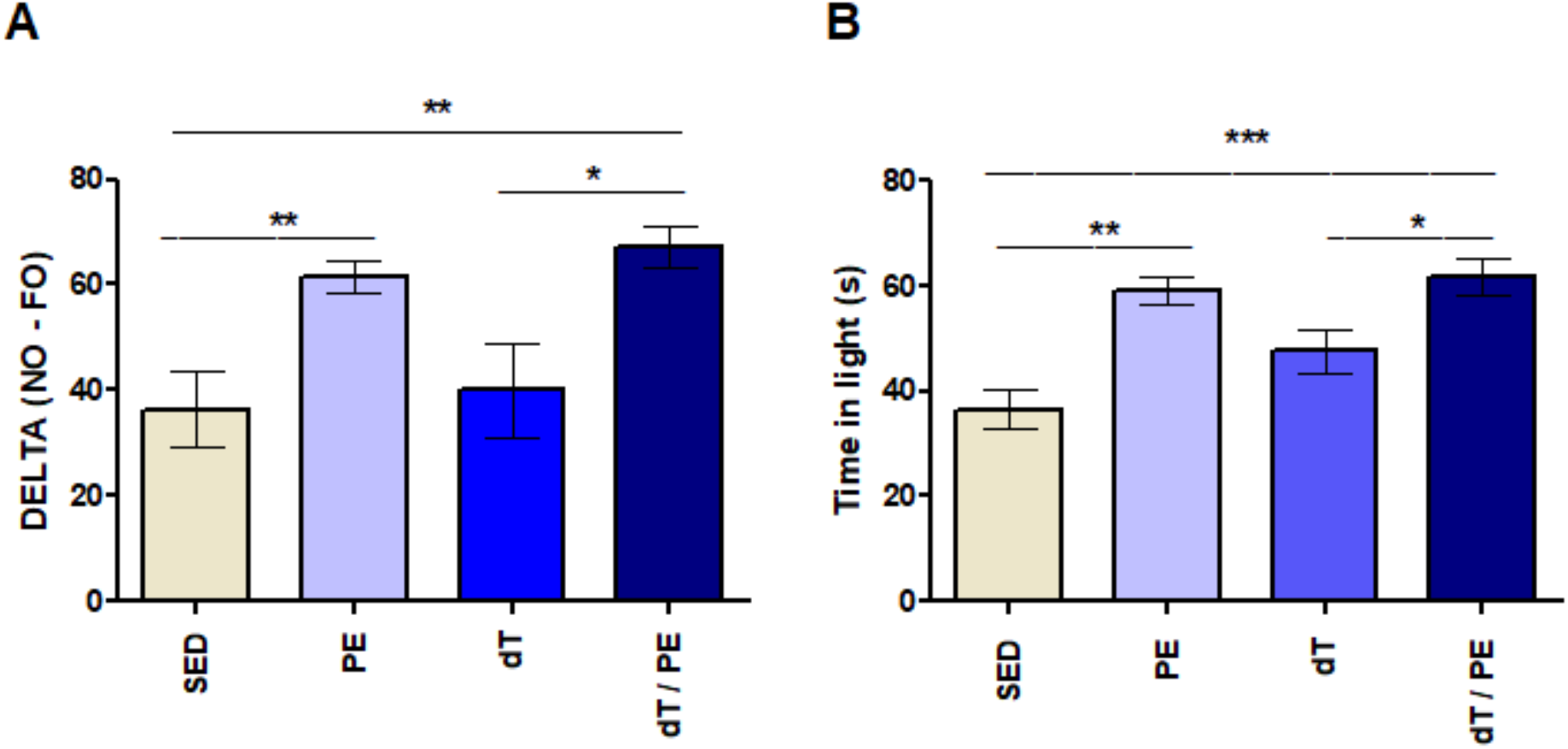
Effect of physical exercise and immunization with Td vaccine. started 14 days after the end of chloroquine treatment, on cognitive (A) and behavioral (B) performance in healthy mice.

Delayed-PE did also improve cognitive parameters, although it did not fully restore parameters to the basal levels of control groups; and also attenuated anxiety-like behavior in infected animals (Figure 4).

**Figure 4.**
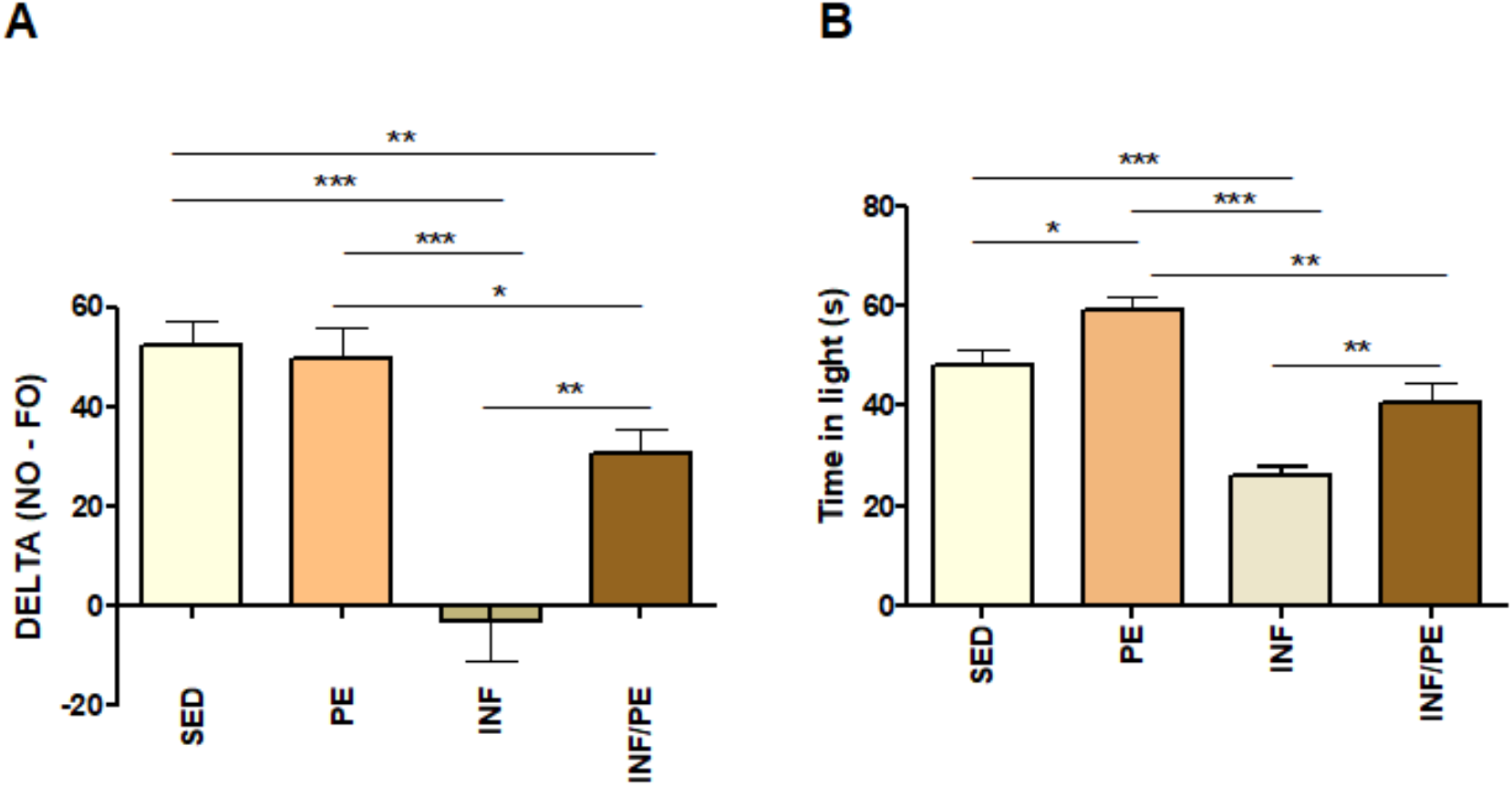
Effect of physical (delayed) exercise. initiated 80 days after the end of treatment with chloroquine, on the attenuation of cognitive (A) and behavioral (B) impairment caused by non-severe murine malaria.

### PRELIMINARY CONCLUSIONS

PE, whether combined with immunization or not, is a promising tool for mitigating cognitive-behavioral sequelae of n-SM. In addition, PE, *per se*, is a potent enhancer of cognitive-behavioral performance in homeostasis, as previously demonstrated in several other reports. Although mild malaria does not require hospitalization, it causes significant neurocognitive impairment, reduces quality of life, and generates economic and health burdens in affected countries. Therefore, studies that aim to evaluate accessible and non-invasive complementary therapeutic strategies - which restore neurocognitive homeostasis lost due to infectious diseases - are increasingly welcome, with malaria as a backdrop, since it is a major public health problem in the world, second only to AIDS and tuberculosis.

## CONFLICTS OF INTERESTS

The authors declare no competing financial interests or personal relationships that could have influenced the opinions expressed in these preliminary results.

## FINANCIAL SUPPORT

LPS (E-26/205.722/2022) receives a Faperj Pos-Doctoral fellowship and IOL receives a Scientific Initiation fellowship from CNPq. CTDR is supported by the *Conselho Nacional de Desenvolvimento Científico e Tecnológico* (*CNPq*, Brazil 310445/2017-5) through a Productivity Research Fellowship, and receives a *Cientista do Nosso Estado* fellowship by the *Fundação Carlos Chagas Filho de Amparo à Pesquisa do Estado do Rio de Janeiro* (*Faperj*, CTDR E-26/202.921/2018). The *Laboratório de Pesquisa em Malária* (LPM-IOC, Fiocruz) is an Associate Laboratory of the *Instituto Nacional de Ciência e Tecnologia em Neuroimunomodulação* of the *CNPq* (*INCT-NIM/CNPq* Project 465489/2014-1) and of the *Rede de Neuroinflamação da Faperj* (*Redes/Faperj*, Project 26010.002418/2019) and receives financial support of the Faperj (Project SEI-260003/001169/2020). Funders did not have any role in paper conceptualization, data interpretation, or writing of the paper.

## Notes

### Competing Interest Statement

The authors have declared no competing interest.

